# Genomic landscape and chronological reconstruction of driver events in multiple myeloma

**DOI:** 10.1101/388611

**Authors:** Francesco Maura, Niccoló Bolli, Nicos Angelopoulos, Kevin J. Dawson, Daniel Leongamornlert, Inigo Martincorena, Thomas J. Mitchell, Anthony Fullam, Santiago Gonzalez, Raphael Szalat, Bernardo Rodriguez-Martin, Mehmet Kemal Samur, Dominik Glodzik, Marco Roncador, Mariateresa Fulciniti, Yu Tzu Tai, Stephane Minvielle, Florence Magrangeas, Philippe Moreau, Paolo Corradini, Kenneth C. Anderson, Jose M. C. Tubio, David C. Wedge, Moritz Gerstung, Herve Avet-Loiseau, Nikhil Munshi, Peter J. Campbell

**Author notes:** These authors contributed equally to this work (Francesco Maura, Niccoló Bolli). Corresponding Authors: Dr Peter J Campbell, Cancer Genome Project, Wellcome Sanger Institute, Hinxton CB10 1SA, United Kingdom., Dr Nikhil C. Munshi Dana-Farber Cancer Institute 450 Brookline Avenue, Dana B106 Boston, MA 02215, USA.

## Abstract

Multiple myeloma (MM) has a heterogeneous genome, evolving through both pre-clinical and post-diagnosis phases. Here, using sequences from 67 MM genomes serially collected from 30 patients together with public datasets, we establish a hierarchy of driver lesions. Point mutations, structural variants and copy number aberrations define at least 7 genomic subgroups of MM, each with distinct sets of co-operating driver mutations. Complex structural events are major drivers of MM, including chromothripsis, chromoplexy and a replication-based mechanism of templated insertions: these typically occur early. Hyperdiploidy also occurs early, with individual chromosomes often gained in more than one chronological epoch of MM evolution, showing a preferred order of acquisition. Positively selected point mutations frequently occur in later phases of disease development, as do structural variants involving *MYC*. Thus, initiating driver events of MM, drawn from a limited repertoire of structural and numerical chromosomal changes, shape preferred trajectories of subsequent evolution.

---

The genome of multiple myeloma (MM) is complex and heterogeneous, with a high frequency of structural variants (SVs) and copy-number abnormalities (CNAs)^1–3^. Translocations between the immunoglobulin heavy chain (*IGH*) locus and recurrent oncogenes are found in ~40% of patients. Cases without *IGH* translocations often have a distinctive pattern of hyperdiploidy affecting odd-numbered chromosomes, where the underlying target genes remain mysterious. These SVs and recurrent CNAs are considered early drivers, being detectable also in pre-malignant stages of the disease^1,2,4^. Cancer genes are also frequently altered by driver point mutations, with MAPK and NF-κB signaling as major targets^5–8^.

Many blood cancers develop along preferred evolutionary trajectories. Early driver events, drawn from a restricted set of possible events, differ in which subsequent cancer genes confer clonal advantage, leading to considerable substructures of co-operativity and mutual exclusivity among cancer genes. These subtypes vary in chemosensitivity and survival, suggesting that although patients share a common histological and clinical phenotype, the underlying biology is distinctly heterogeneous. Preliminary studies have suggested that these patterns exist in MM as well^5,7,9–12^, but have not yet been systematically defined in large cohorts with broad sequencing coverage.

We performed whole genome sequencing (WGS) of 67 tumor samples collected at different time points from 30 MM patients, together with matched germline controls (**Supplementary Fig. 1**, **Supplementary Table 1, Methods**). We also included in our analyses published whole exome data from 804 patients^13,14^. To discover significant cancer genes, we analyzed the ratio of non-synonymous to synonymous mutations, correcting for mutational spectrum and covariates of mutation density across the genome using a published algorithm^15,16^. Overall, 55 genes were significantly mutated with a false discovery rate of 1% (Fig. 1a, **Supplementary Table 2**). A significant fraction of these driver mutations was detected at subclonal level, suggesting a major role in late phases of cancer development (**Supplementary Fig. 2a**). Beyond well-known myeloma genes such as *KRAS*, *NRAS*, *DIS3* and *FAM46C*^5–8^, several other interesting candidate genes emerged. The linker histones *HIST1H1B*, *HIST1H1D, HIST1H1E* and *HIST1H2BK* all showed a distinctive pattern of missense mutations clustered in the highly conserved globular domain (**Supplementary Figure 2b-e**), as reported in follicular lymphoma^17^. Many of the mutations were nearby, or directly affected, conserved positively charged residues critical for nucleosome binding, suggesting that they disrupt the histones’ role in regulating higher order chromatin structure. *FUBP1*, an important regulator of *MYC* transcription^18^, showed an excess of splice site and nonsense mutations, suggesting it may be a tumor suppressor gene in MM (**Supplementary Figure 2f**). *MAX*, a DNA-binding partner of *MYC*, showed an interesting pattern of start-lost mutations, nonsense and splice site mutations, together with hotspot missense mutations at residues Arg35, Arg36 and Arg60, known to abrogate DNA binding^7^ (**Supplementary Fig. 2g**). Genes with rather more mysterious function were also significant: the zinc finger *ZNF292*, recently described as mutated in chronic lymphocytic leukemia and diffuse large B-cell lymphoma^19,20^, showed an excess of protein-truncating variants (**Supplementary Fig. 2h**); the uncharacterized *TBC1D29* gene showed two hotspots of missense mutations^21^ (**Supplementary Fig. 2i**).

**Figure 1.**
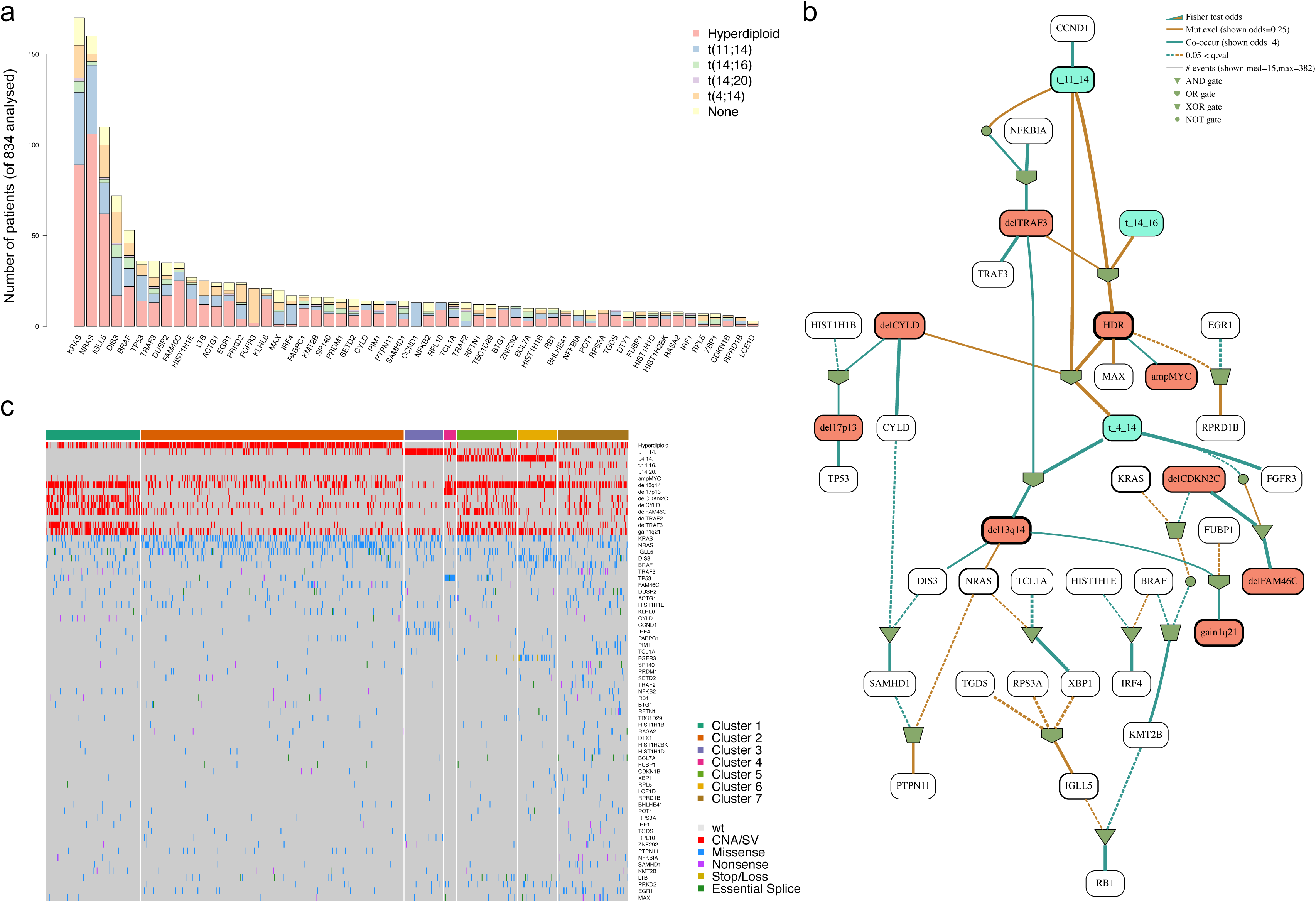
a) Landscape of Driver Mutations in Multiple Myeloma (MM). Each bar represents a distinct driver gene and each bar’s colour indicate its prevalence across the main MM cytogenetic sub-groups. b) We built the optimal Bayesian network by considering the recurrent SVs/ CNAs (n 14) and driver SNVs (n 55) across 724 MM patients where the final list of 69 variables was assessed. To further investigate the type of recurrence patterns we fitted logic gates between parent and child nodes in the network. The gate combination with the highest Fisher exact test p value was selected. The line width is proportional to the log hazard ratio of the test. Dashed lines represent non-significant associations (p>0.05). CNAs and translocations were coloured by red and light blue respectively. The width of the boundary line of each drawn box is proportional to its prevalence across the entire series. c) Heat map showing the main MM genomic subgroups across 724 MM patients. The genomic profile of each cluster was generated by integrating the hierarchical Dirichlet process and Bayesian network data. Rows in the graph represent individual genomic lesions, and the columns represent patients.

**Figure 2.**
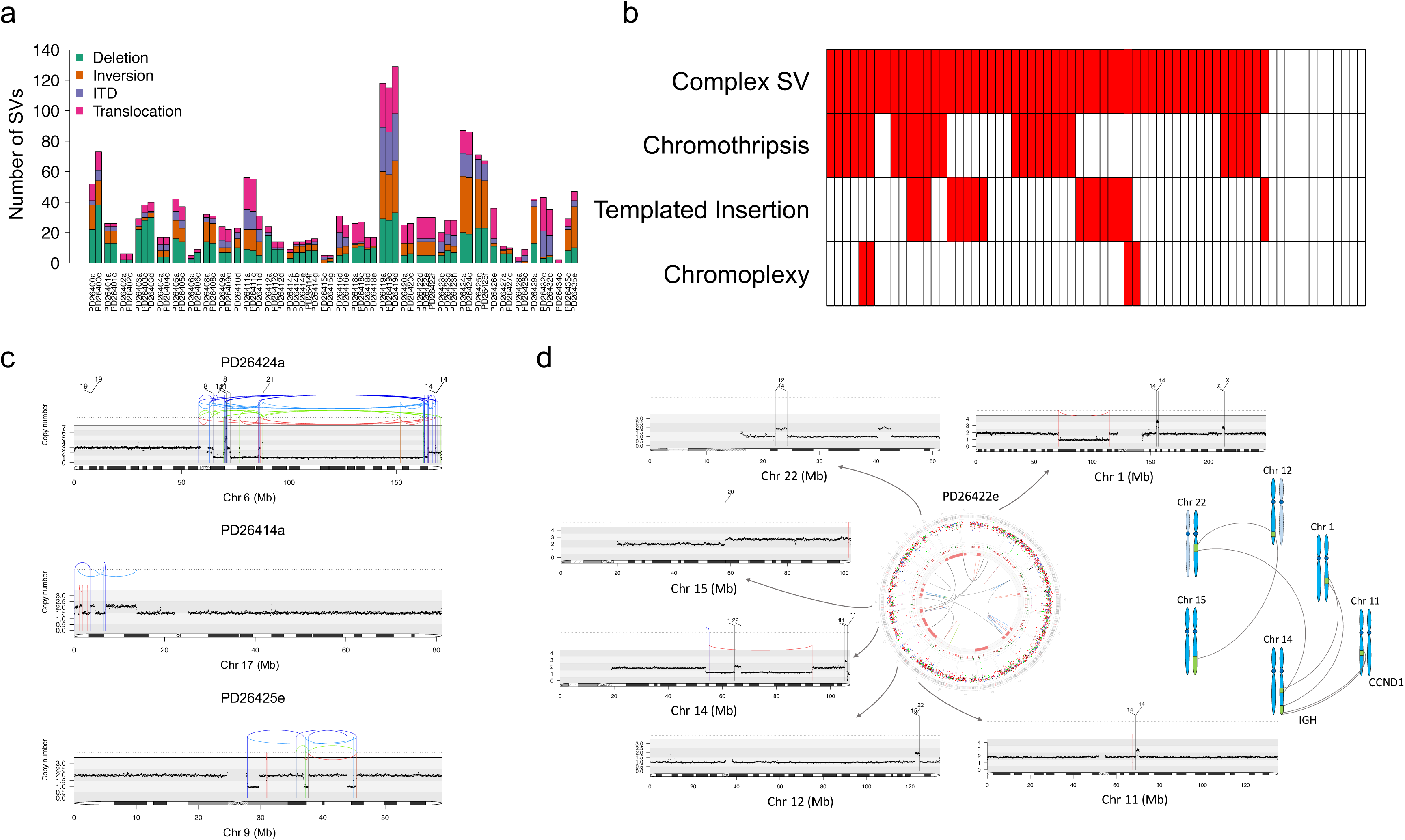
a) SVs prevalence across the entire series. b) Heatmap representing the distribution and prevalence of the main complex SVs: chromothripsis, chromoplexy and templated insertion. c) Three examples of chromothripsis. d) Example of templated insertion. In the middle, the genome plot of patient PD26422 represents all main genomic events: mutations (external circle), indels (middle circle; dark green and brown lines represent insertion and deletion respectively), copy number variants (red = deletions, green = gain) and rearrangements (blue = inversion, red = deletions, green = ITD, black = translocation). Externally, a copy number/rearrangement plot of each chromosome involved by the templated insertion is provided, highlighting a focal CNA around each breakpoint. This case represents a clear example of how templated insertion may involve critical driver oncogenes, like *CCND1* in this case. A schematic representation of this sample templated insertion is reported on the right.

In pairwise comparisons, these cancer genes showed distinct patterns of co¬mutation and mutual exclusivity (**Supplementary Fig. 2j**). To define the logic rules underpinning the conditional dependencies of driver events, we employed Bayesian networks and the hierarchical Dirichlet process (Fig. 1b-c, **Supplementary Fig. 3**). This confirmed that the strongest determinants of genomic substructure in myeloma are *IGH* translocations and recurrent CNAs. Some co-operating genetic lesions were non-randomly distributed across these main groups: a significant fraction of patients without *IGH* translocations were generally enriched for 1q gain and deletions on 1p13, 1p32, 13q, *TRAF3* and *CYLD* (*Cluster 1*). RAS signaling mutations, especially *NRAS* and *KRAS*, were associated with hyperdiploidy and *MYC* translocations (*Cluster 2*). A significant fraction (33%) of patients with t(11;14) were characterized by low genomic complexity and high prevalence of *IRF4* and *CCDN1* mutations (*Cluster 3*). Patients harboring *TP53* bi-allelic inactivation were clustered in an independent sub group (*Cluster 4*). A significant fraction of *MMSET* (51%) and *CCND1* (19%) translocated patients were characterized by multiple cytogenetic aberrations, with low prevalence of *CYLD/FAM46C* deletions and *TRAF3* deletion respectively *(Cluster 5).* A second fraction of patients with *MMSET* translocation (46%) were grouped with deletion of 13q14, gain of 1q21, *DIS3* and *FGFR3* mutations (*Cluster 6*); finally, patients with either *MAF*/*MAFB* translocations or no *IGH* translocations was characterized by a high driver mutation rate (*Cluster 7*). Thus, there is evidence for at least 7 distinct genomic subtypes of myeloma, each with distinct combinations of driver mutations and recurrent SVs (Figure 1c).

**Figure 3.**
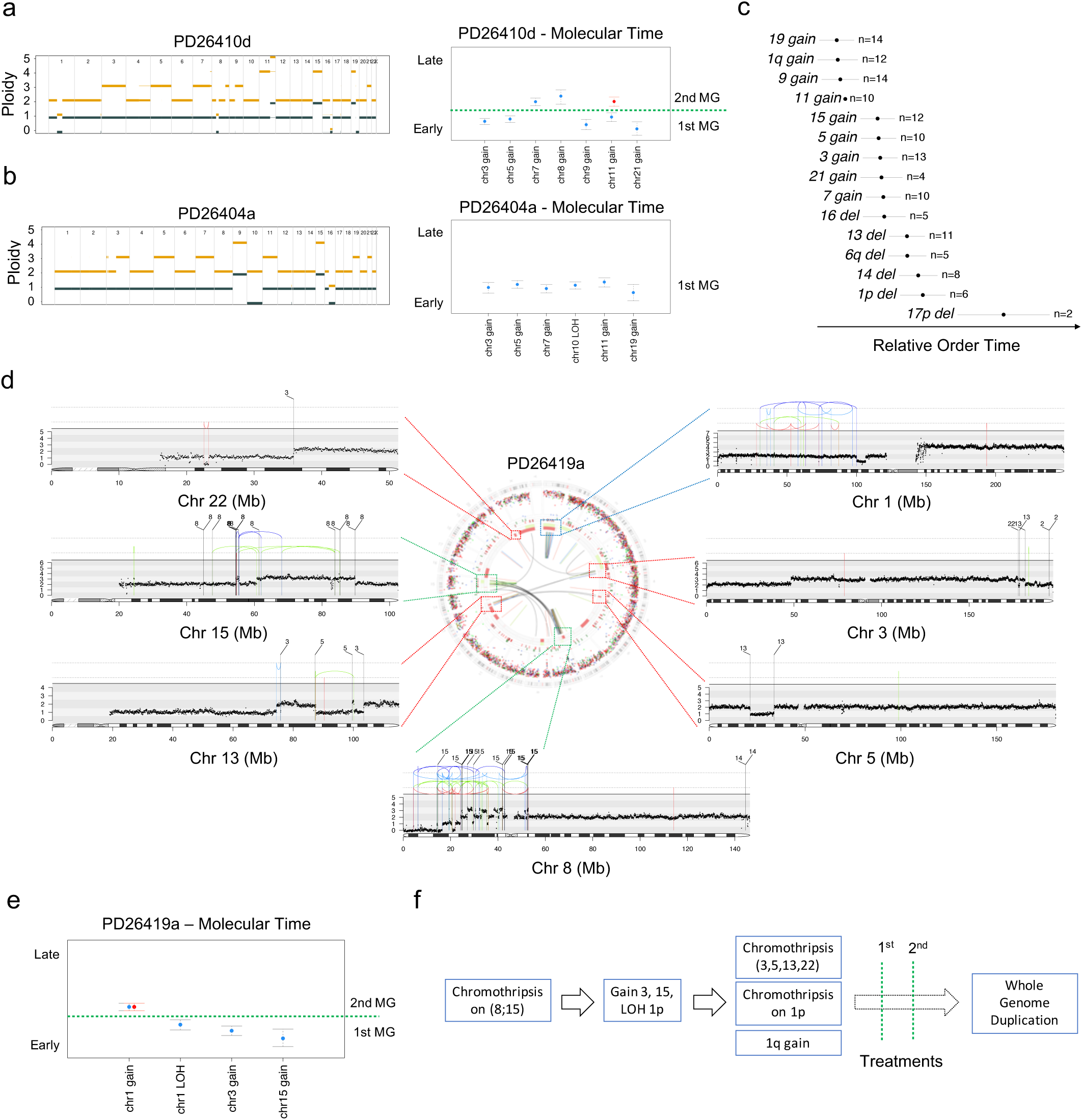
a-b) Molecular time estimation on all chromosomal gains observed in 2 hyperdiploid MMs. On the left the copy number profile is reported, where gold line = total copy number, grey = copy number of the minor allele. The presence of more than 1 cytogenetic segment is compatible with the existence of a subclonal CNA whose CCF is proportional to the segment thickness [see for example the gain on 4q in PD26410d (a)]. On the right, the molecular time (blue dots) estimated for each clonal gain and copy neutral loss of heterozygosity (**Methods**). Red dots represent the molecular time of a second extra gain occurred on a previous one. Dashed green lines separate multi gain events occurring at different time windows c) Bradley Terry model based on the integration between the CCF and molecular time of each recurrent MM CNAs (gains and deletions). Segments were ordered from the earliest (top) to the latest (bottom) occurring in relative time from sampling (X-axis). d) Genome plot of patient PD26419a where the three chromothripsis events [(8;15), (3;5;13;22) and 1p] were highlighted with different colored dashed lines connected to specific rearrangements/copy number plots. In these plots, the red arch represents a deletion, the green arch represents an ITD and the blue arch represents an inversion. e) Molecular time of the main clonal gains and LOH in the PD26419a sample. This data suggested the existence of at least 2 different and independent time windows: the first involving alterations on chromosome 3, 15 and 1p and the second on chromosome 1q. f) The driver events of patient PD26419 are reconstructed in chronological order.

Straightforward reciprocal translocations such as the canonical *IGH*-oncogene translocations only accounted for 6% (127/2113) of SVs in the 67 samples from 30 patients studied by WGS (Figure 2a, **Supplementary Fig. 4a-b**). Other structural variants included many unbalanced translocations and complex events (**Supplementary Fig. 4c-f, Methods**)^22^. Most (24/30; 80%) patients had at least one complex SV, of which chromothripsis was the most frequent (11/30; 36%) (Fig. 2b-c)^22–26^. Chromoplexy occurred in 3 patients (Fig. 2b, **Supplementary Fig. 5a-c**)^22,27^.

**Figure 4.**
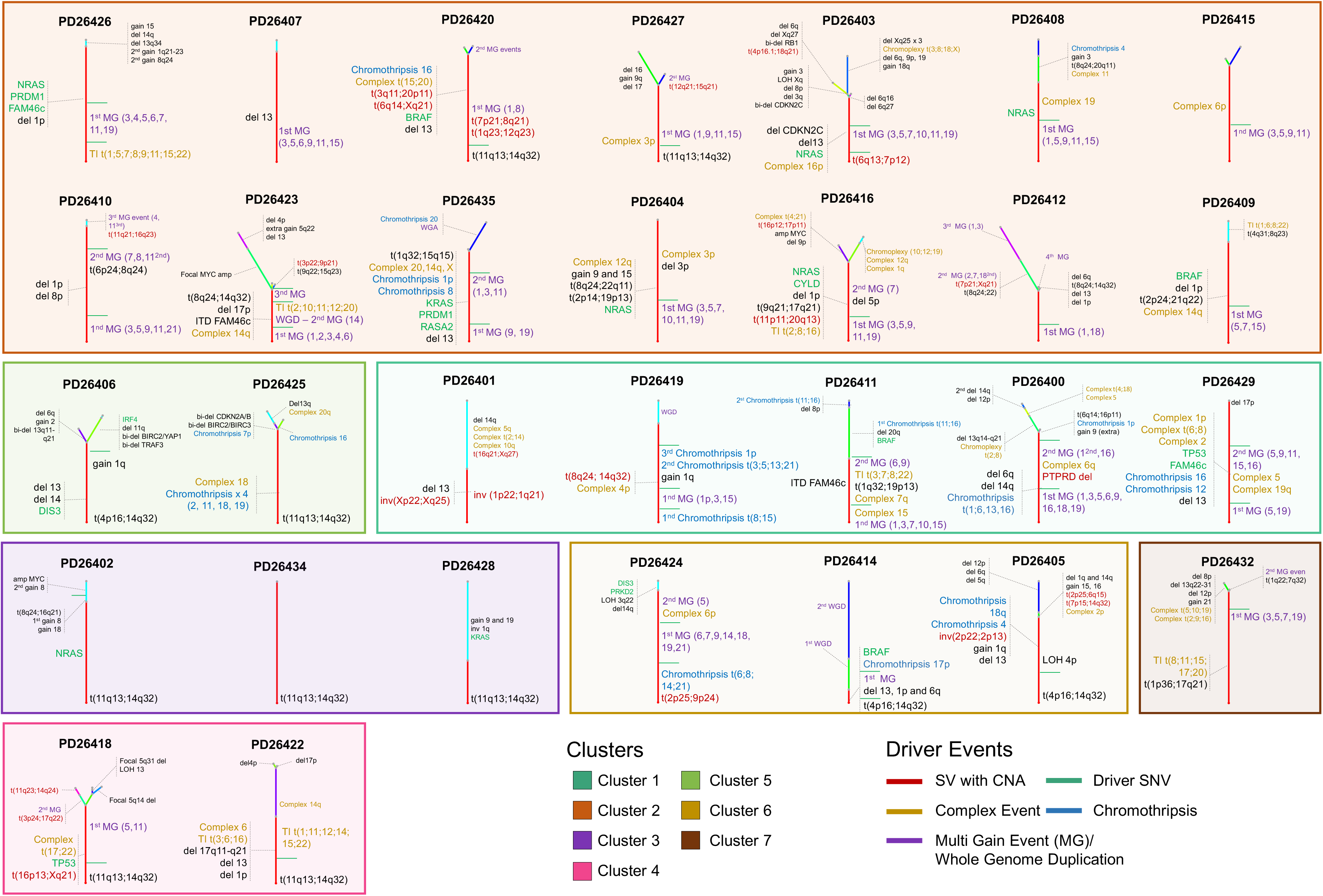
The most likely phylogenetic trees generated from the Dirichlet process analysis (**Methods**). The root (always colored in red) and branch length is proportional to the (sub)clone mutational load. All main drivers (CNAs, SNVs and SVs) were annotated according to their chronological occurrence. Early clonal events (root), where it was possible to establish a specific time window, were chronologically annotated on the right. All different “root” time windows were separated by a green line; conversely, early drivers without a clear timing were grouped together on the left of the root. All driver events that occurred in the root were reported with larger font size. Patients were grouped according to the genomic clustering showed in Figure 1c. Templated insertion is abbreviated with TI.

In 6/30 (20%) patients, we found a novel complex pattern characterized by cycles of templated insertions. Here, several low-amplitude copy number gains on different chromosomes were linked together through SVs demarcating the region of duplication (Fig. 2b,d; **Supplementary Fig. 5d-h**). The most plausible explanation for this pattern is that the templates are strung together into a single chain, hosted within one of the chromosomes. We have observed such events in some solid cancers^22^, where the copy number gains suggest a mutational process that is replication-based, rather than the break-and-ligate processes generating chromothripsis and chromoplexy^23^.

This complex landscape of SVs in myeloma often involved known driver genes: *MYC* (14/30 cases; 46%), *CCND1* (7/30; 23%) and *MMSET* (3/30; 10%) were common targets (**Supplementary Fig. 4b**) of these non-canonical events. The juxtaposition of *CCND1* to the *IGH* locus was caused by either unbalanced translocations or insertional events in 5/8 patients (**Supplementary Fig. 6**). Similarly, *MYC* translocations showed unanticipated complexity, with four cases of templated insertions involving MYC or its regulatory regions (**Supplementary Fig. 7**). Such events are the structural basis of oncogene amplification observed by FISH in many cases of t(11;14) and t(8;14)^28,29^. Interestingly, many of the *MYC* SVs involved the immunoglobulin light chain loci, *IGK* or *IGL*, rather than the heavy chain *IGH* locus, and were seen in patients with hyperdiploidy (**Supplementary Fig. 4b**). Although sometimes occurring late, these events were under strong selective pressure: we identified a striking case of convergent evolution where a subclone bearing an *IGL*:*MYC* translocation was lost and one bearing an *IGH*:*MYC* was acquired at relapse (**Supplementary Fig. 8**). SVs also led to loss of tumor suppressor genes such as *BIRC2/3*, *CDKN2A/B, CDKN2C*, *TRAF3*^30^ and *FAM46C*, either within focal deletions or more complex events. These data confirm that SVs, accessing both simple and complex mechanisms of genome rearrangement, is a major force shaping the myeloma genome.

In our cohort of serial WGS samples we could evaluate the temporal evolution of the disease, integrating SVs, CNAs and point mutations. This included not only defining clonal and subclonal mutations and CNAs, but also timing the relative order of acquisition of clonal chromosomal gains, which we estimated with “molecular time” analysis based on the fraction of duplicated to single-copy point mutations (**Methods**)^31^.

Hyperdiploidy was found in 18/30 patients. Applying our “molecular time” analysis, we found that chromosomal gains were not necessarily acquired simultaneously (Fig. 3a). In 11/18 cases, we found evidence of several independent gains over time, although the other 7 patients acquired all chromosome gains in a much narrower time window (Fig. 3b). As might be expected if gains occur as independent events, subclonal evolution within the myeloma cells between diagnosis and relapse could lead to considerable diversification of chromosome complements. In fact, we observed significant karyotype changes within the same patient over time, including loss of some trisomies in hyperdiploid patients (**Supplementary Fig. 9a-b)**^32^. At the extreme end of this cytogenetic dynamism, we found 4 patients acquiring a whole genome duplication at relapse, suggesting its potential role in relapsed/refractory stages (**Supplementary Fig. 9c-g**).

By pairwise comparison of the relative timings of copy number alterations we reconstructed the preferred chronological order of CNAs acquisition (**Methods**). Gains of chromosomes 19, 11, 9 and 1q were amongst the earliest in our series (Fig. 3c) and recurrent chromosome losses were generally acquired later than trisomies, consistent with the proposition that hyperdiploidy is an early driver event in MM.

Translocations involving *CCND1* and *MMSET* were always fully clonal, similarly confirming their early driver role in MM pathogenesis. Most chromothripsis events were clonal and conserved during evolution (17/22; 77%), suggesting they occurred early in MM pathogenesis. However, a small fraction of patients showed some evidence of subclonal or late chromothripsis (5/22; 23%), implying a potential involvement in drug resistance and late cancer progression (**Supplementary Fig. 9h**).

We integrated all extracted chronological data on SVs, hyperdiploidy and point mutations to generate phylogenetic trees for each sample (**Methods** and **Supplementary Fig. 10**)^5,33^. The methodology is worked through for one illustrative patient carrying i) several chromosome gains, ii) 3 separate chromothripsis events and iii) a whole genome duplication (Fig. 3d-f). One chromothripsis involved chromosomes 8 and 15, duplicating the long arm of chromosome 15. Because few mutations were present on chromosome 15 at the time it duplicated, this must have occurred early in molecular time (Fig. 3e). Gain of chromosome 3 and copy-neutral loss-of-heterozygosity of small arm of chromosome 1 (chr1p) occurred not long after, and were followed by a second chromosomal crisis involving chromosomes 3, 5, 13 and 22. This chromothripsis must have occurred in one of the two duplicated alleles of chromosome 3 (and therefore after the acquisition of a chromosome 3 trisomy) because losses within the chromothripsis region had copy number 2 and SNPs were heterozygous. Within the same time window, a separate chromothripsis event occurred on chr1p after copy-neutral loss-of-heterozygosity. Finally, this patient underwent whole genome duplication after two therapy lines (**Supplementary Fig. 9f**).

Applying this approach to all patients, we observed that the trunks of the phylogenetic trees of 29/30 (97%) patients were characterized by few genomic events generally acquired during different time windows of the MM life history before emergence of the most recent common ancestor (Fig. 4). Overall, chromothripsis, cycles of templated insertions, chromosomal gains and other SVs accounted for most of the earliest events, emerging as key early drivers of the disease and paving the way for subsequent driver mutations that would confer further selective advantage to the clone.

Taken together, these data suggest that MM development follows preferred evolutionary trajectories, with stuttering accumulation of driver events in keeping with its insidiously progressive but unpredictable clinical course. Critical early events include immunoglobulin translocation with *MMSET* and *CCND1*; hyperdiploidy and focal complex structural variation processes hitting key myeloma genes. These early driver mutations shape the subsequent evolution of myeloma, each with preferred sets of co-operating cancer genes.

## Acknowledgements

FM is supported by A.I.L. (Associazione Italiana Contro le Leucemie-Linfomi e Mieloma ONLUS) and by S.I.E.S. (Società Italiana di Ematologia Sperimentale).

NB is funded by AIRC (Associazione Italiana per la Ricerca sul Cancro) through aMFAG (n.17658).

This work was supported by: Department of Veterans Affairs Merit Review Award I01BX001584-01 (NCM), NIH grants P01-155258 (N.C.M., H.A.L., M.F., P.J.C., K. C.A.) and 5P50CA100707-13 (N.C.M., H.A.L., K.C.A).

## Authorship Contributions

F.M., N.B., and P.J.C. designed the study, collected and analyzed the data and wrote the paper; H.A.L., N.C.M. designed the study and collected the data; K.J.D., D.L., N.A., I.M., T.J.M., A.F., D.G., S.G., M.G., M.R., F.A., B.R.M., J.M.C.T. and D.W. analyzed the data; S.M., R.S., M.K.S., M.F., Y.T.T., M.F., P.M., P.C., K.C.A., collected the data.

